# Impact of COVID-19: Decrease in the Number of Fledging Barn Swallow Chicks in Tokyo

**DOI:** 10.1101/2020.11.16.380899

**Authors:** Takuma Hayashi, Nobuo Yaegashi, Ikuo Konishi

**Affiliations:** National Hospital Organization Kyoto Medical Center, Kyoto, Japan; START, Japan Science and Technology Agency, Tokyo, Japan; Department of Obstetrics and Gynecology, Tohoku University School of Medicine, Miyagi, Japan; Department of Obstetrics and Gynecology, Kyoto University School of Medicine, Kyoto, Japan; Immediate Past President, Asian Society of Gynecologic Oncology, Tokyo, Japan

**Author notes:** Corresponding Author, Takuma Hayashi, DMSci, GMRC, MS, PhD, National Hospital Organization, Kyoto Medical Center Fukakusa Mukai-Cho, Fushimi-Ku, Kyoto-city, Kyoto, Japan.

**Keywords:** COVID-19, barn swallow, nest, 23 districts of Tokyo

## Abstract

Barn swallows that have crossed the sea from Southeast Asia usually appear in the Kyushu Region of Japan around March after passing through Okinawa Prefecture. When the climate becomes warmer, these birds then move further north, nesting and raising their chicks in various parts of Japan. It is worth noting that barn swallows typically nest on man-made objects, for example, the roofs of houses and barns. It is believed that this is because barn swallows protect their eggs and chicks from foreign enemies such as sparrows and crows so they build their nests in populated areas. The barn swallow’s behavior of using the presence of people to keep foreign enemies away shows that barn swallows are quite wise. However, it has been reported that from the spring to summer of 2020, barn swallows, nesting and raising their chicks, which were seen every year, were not found in various parts of Japan. Therefore, we investigated the relationship between people’s self-restraint from going out and the fledging of barn swallow chicks in Tokyo metropolitan during the corona virus disease 2019 (COVID-19) era. The results of the survey showed a link between people’s refraining from going out and the fledging of barn swallow chicks. Next spring of 2021, the termination of COVID-19 is an important environment for swallow chick fledging.

## Introduction

Barn swallows breed in wide areas of the Northern Hemisphere.^1,2^ In Japan, they also breed in Okinawa and other prefectures in the north. The main wintering locations of barn swallows that breed in Japan are Taiwan, the Philippines, northern Borneo, the Malay Peninsula, and Java. The total length of a barn swallow is approximately 17 cm, and its wingspan is approximately 32 cm.^3^ Its wings are large, and the body is elongated to suit flight. In addition, its legs are short and unsuitable for walking; thus, these birds rarely land on the ground except when seeking nesting mud.

Barn swallows build nests by solidifying mud and dead grass with saliva. In Japanese cities, their nests are mostly built on man-made objects. They tend to breed in the same places where people live, such as the eaves of private houses. It is thought that natural enemies such as crows and sparrows are difficult to approach as the reason why barn swallows behave like this action.^(Note 1)^ Sparrows are considered as birds that build nests in private houses. However, the behavior of barn swallows is very different from that of sparrows and other birds in terms of building a nest in populated areas.

Barn swallows build their nests in a conspicuous place to protect their nests, eggs, and chicks from the attacks of other birds and snakes. It is believed that barn swallows use humans as a security guard. Sparrows that rob their nests and crows that aim for eggs and chicks do not approach places where people are always present or where people come and go. Weak barn swallows are thought to cover their weakness by nesting near humans.

Barn swallows usually build new nests, but they sometimes repair old nests and reuse them. Their breeding season is from April to July. One female barn swallow lays three to seven eggs and mainly warms them. Barn swallows have 13–17 days of egg-bearing and 20–24 days of brooding in their nests. The fledging rate of the first breeding of barn swallows is estimated to be 50%. The same pair of barn swallows, whether successful or unsuccessful in the first breed, then breeds the second time.

The barn swallows’ nest building at a convenience store near Dogo Onsen, the oldest hot spring in Japan, has become a hot topic. Every spring, these birds nest on the signboards of convenience stores. Neighboring residents are supporting the fledging of swallows.

Around May 2020, barn swallows built their nest on the signboard of a convenience store. These birds laid four eggs; however, they left their eggs in place and flew away. This may be because there were no people in the city due to the outbreak of COVID-19. Barn swallows seemed to have noticed that people refrain from going out. Barn swallows seem to be protected from foreign enemies such as crows by people who are going out in the city.

## Materials and Methods

It has been pointed out that in the COVID-19 era, barn swallows may not nest and raise their chicks as usual because people are refraining from going out. Therefore, we examined the situation of the swallows’ nesting and fledging of chicks in Tokyo metropolitan during this era. As a concrete research method, using the material^4^ of the National Survey of Barn Swallows of the Wild Bird Society of Japan, we measured the number of barn swallow chicks that flew out of the nest in the entire 23 districts of Tokyo metropolitan and the area within the Yamanote Line in each year from 2015 to 2020.

In addition, using the document analyzed by Agoop Corporation (Tokyo, Japan),^5^ we compared and considered the number of people who went out during the daytime in the entire 23 districts of Tokyo metropolitan and the area within the Yamanote Line in July 2019^(Note 2)^ and who went out during the daytime in the same areas during the outbreak of COVID-19 in July 2020. From these analysis results, we examined the relationship between the number of people who went out and that of barn swallow chicks that flew out of the nest.

In the COVID-19 era, swallow investigators are observing and investigating the paired barn swallows’ nests and chicks flying out of their nests once a day in their area of responsibility. Therefore, the investigation of the swallows has not been affected by people’s refraining from going out due to COVID-19.

This study was reviewed and approved by the Central Ethics Review Board of the National Hospital Organization Headquarters in Tokyo, Japan. The authors attended a 2020 educational lecture on medical ethics supervised by the Japanese government. The completion numbers of the authors are AP0000151756, AP0000151757, AP0000151769, and AP000351128.

The Global Positioning System function of mobile phones was used to measure the number of people who came and went around the stations of Japan Railway Companies and core facilities in major cities in Japan. Therefore, this study did not require the consent of the participants.

## Results

We compared the outing rate of people in the 23 districts of Tokyo metropolitan in July 2020 with that of those in the same area during the same period in 2019. The areas within the Yamanote Line within the entire 23 districts of Tokyo metropolitan, near Shibuya Station, Omotesando Station, Harajuku Station, Shinjuku Station, Shinjuku Kabukicho, Ikebukuro Station, Akihabara Station, Tokyo Station, Shimbashi Station, Shinjuku Station, Ueno Station, and Roppongi Station, were surveyed.

The results of the survey showed that the outing rates in the 23 districts of Tokyo metropolitan were as follows: −21.7% near Shibuya Station, −30.0% near Shibuya Center Street, −17.50% near Omotesando Station, −27.80% near Harajuku Station, −30.30% near Shinjuku Station, −29.40% near Shinjuku Kabukicho, −18.30% near Ikebukuro Station, −24.50% near Akihabara Station, −35.00% near Tokyo Station, −36.13% near Shimbashi Station, −42.10% near Shinagawa Station, −33.80% near Ueno Station, −22.40% near Asakusa Station, −34.50% near Ginza Station, −39.80% near Roppongi Station, −49.70% near Haneda International Airport Terminal Station, and −45.60% in Odaiba.

In 2020, there were 120 fledging barn swallow chicks in all 23 districts of Tokyo metropolitan and 26 in the area within the Yamanote Line, with a ratio of 26% (26/120). In 2019, there were 107 fledging barn swallow chicks in all 23 districts of Tokyo metropolitan and 42 in the area within the Yamanote Line, with a ratio of 39% (42/107). In 2018, there were 205 fledging barn swallow chicks in all 23 districts of Tokyo metropolitan and 80 in the area within the Yamanote Line, with a ratio of 39% (80/205). In 2017, there were 250 fledging barn swallow chicks in all 23 districts of Tokyo metropolitan and 77 in the area within the Yamanote Line, with a ratio of 32% (77/250). In 2016, there were 188 fledging barn swallow chicks in all 23 districts of Tokyo metropolitan and 68 in the area within the Yamanote Line, with a ratio of 36% (68/188). In 2015, there were 219 fledging barn swallow chicks in all 23 districts of Tokyo metropolitan and 79 in the area within the Yamanote Line, with a ratio of 36% (79/219).

These research results demonstrated that the ratio of the number of fledging barn swallow chicks in the area within the Yamanote Line to that of fledging swallow chicks in the entire 23 districts of Tokyo metropolitan was extremely low in July 2020, the COVID-19 era, than those in the previous years.

In the COVID-19 era, swallow investigators are observing and investigating the paired barn swallows’ nests and chicks flying out of their nests once a day in their area of responsibility. Therefore, the investigation of the swallows has not been affected by people’s refraining from going out due to COVID-19.

## Discussion

Due to barn swallows’ habit of “building a nest in the environment where people live,” these birds’ nest building is a reference for houses with several people coming and going and houses with prosperous business in local cities in Japan. Therefore, their nest building is a sign of a prosperous business. There is also a legend that a house with a barn swallow’s nest is safe. As such, after their chicks have left the nest, the landlord often carefully leaves the swallow’s nest. In this way, since ancient times, barn swallows and Japanese life have been closely related.

Every year, a pair of barn swallows builds a nest at the eaves of a house, and the chicks leave the nest. However, in the spring of 2020, it has been reported that a pair of barn swallows nested at the eaves of a house and laid eggs but flew away without warming them.^6,7^ Alternatively, there have been reports of barn swallows not flying at all.

Experts from the Wild Bird Society of Japan report that barn swallows have a habit of breeding near humans to protect themselves from foreign enemies such as crows. Barn swallows do not nest in places where the population decreases during the day, such as in residential areas. Instead, they build their nests in places where they see a lot of people. The reason why barn swallows disappeared from the eaves of convenience stores may be the decrease in the number of people going out due to the outbreak of COVID-19.

In October 2020, the outbreak of COVID-19 occurred in Western countries such as the United States, the United Kingdom, France, Spain, and Belgium, and the blockades of major cities in each country were being carried out again. The fight between COVID-19 and humankind will continue until a vaccine against COVID-19 is developed and becomes available worldwide. If, in the spring of 2021, we can see a pair of swallows nesting at the eaves of a house and swallow chicks flying out of the nest in Tokyo metropolitan, in July 2021. The Tokyo Olympics and Paralympics will be held, and people all over the world will sincerely cheer for the brave appearance of the athletes.

## Footnotes

Note 1 On rare occasions, a large number of breeding pairs can be seen nesting on the ceiling of a self-employed private house.

Note 2 The breeding season of barn swallows is from April to July.

## Data Sharing

Data are available on various websites and have been made publicly available (more information can be found in the first paragraph of the “Results” section).

## Acknowledgments

We thank Professor Richard A. Young (Whitehead Institute for Biomedical Research, Massachusetts Institute of Technology, Cambridge, MA) for his research assistance. This study was supported in part by grants from the Japan Ministry of Education, Culture, Sports, Science and Technology (No. 24592510, No. 15K1079, and No. 19K09840), Foundation of Osaka Cancer Research, Ichiro Kanehara Foundation for the Promotion of Medical Science and Medical Care, Foundation for Promotion of Cancer Research, Kanzawa Medical Research Foundation, Shinshu Medical Foundation, and Takeda Foundation for Medical Science.

## Author Contributions

T.H. performed most of the experiments and coordinated the project; T.H. and N.Y. conceived the study and wrote the manuscript. N.Y. and I.K. gave information on clinical medicine and oversaw the entire study.

## Transparency Document

The transparency document associated with this article can be found in the online version at http://.

---

**Figure 1.**
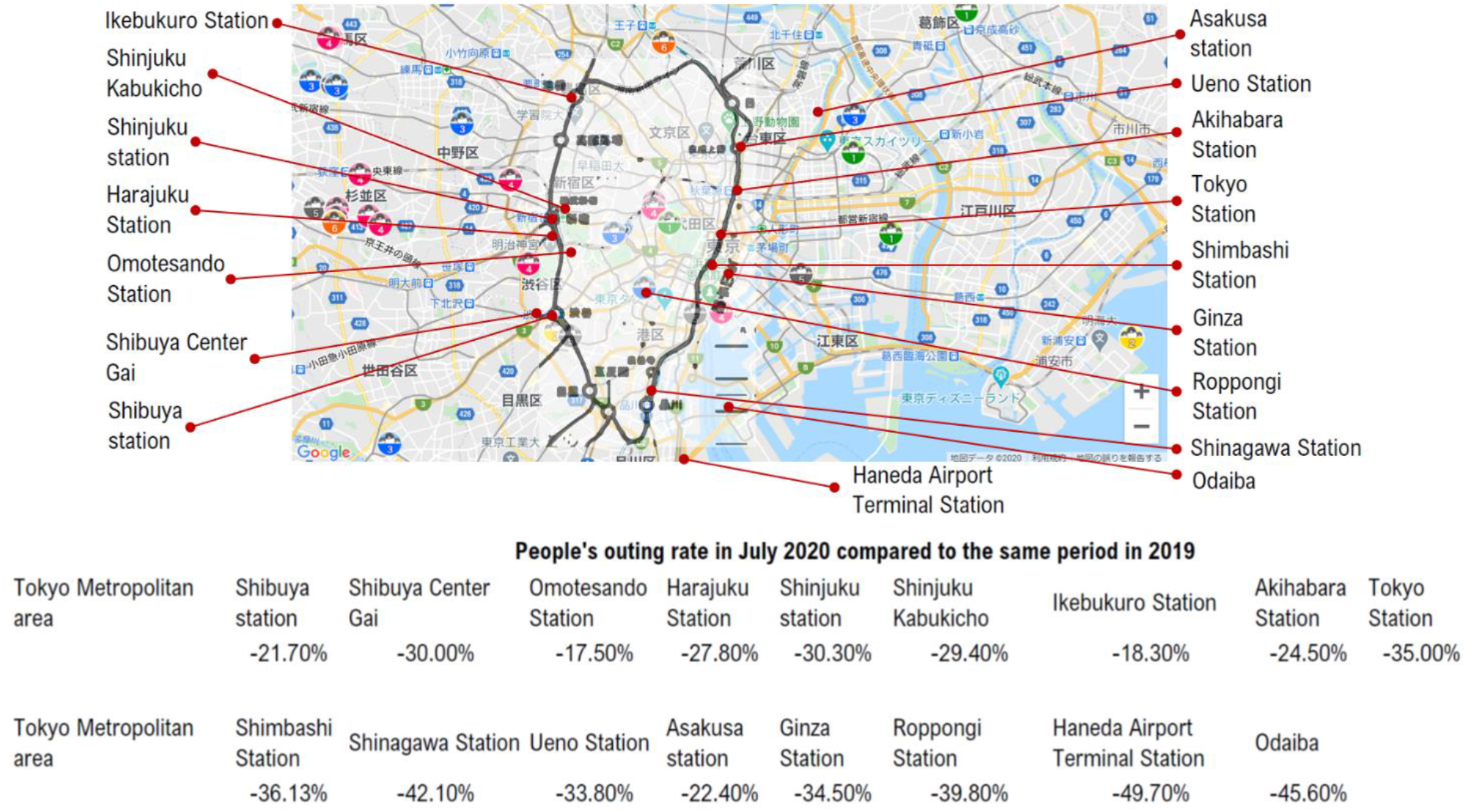
The number of people going out during the daytime in the entire 23 districts of Tokyo metropolitan in the COVID-19 era. The document analyzed by Agoop Corporation (Tokyo, Japan)^5^ reveals the number of people who went out during the daytime in the entire 23 districts of Tokyo metropolitan and the area within the Yamanote Line in July 2019^(Note 2)^ and who went out during the daytime in the same areas during the outbreak of COVID-19 in July 2020. The number of people who went out during the daytime in the entire 23 districts of Tokyo metropolitan and the area within the Yamanote Line during the outbreak of COVID-19 in July 2020 is significantly lower than that in July 2019.

**Figure 2.**
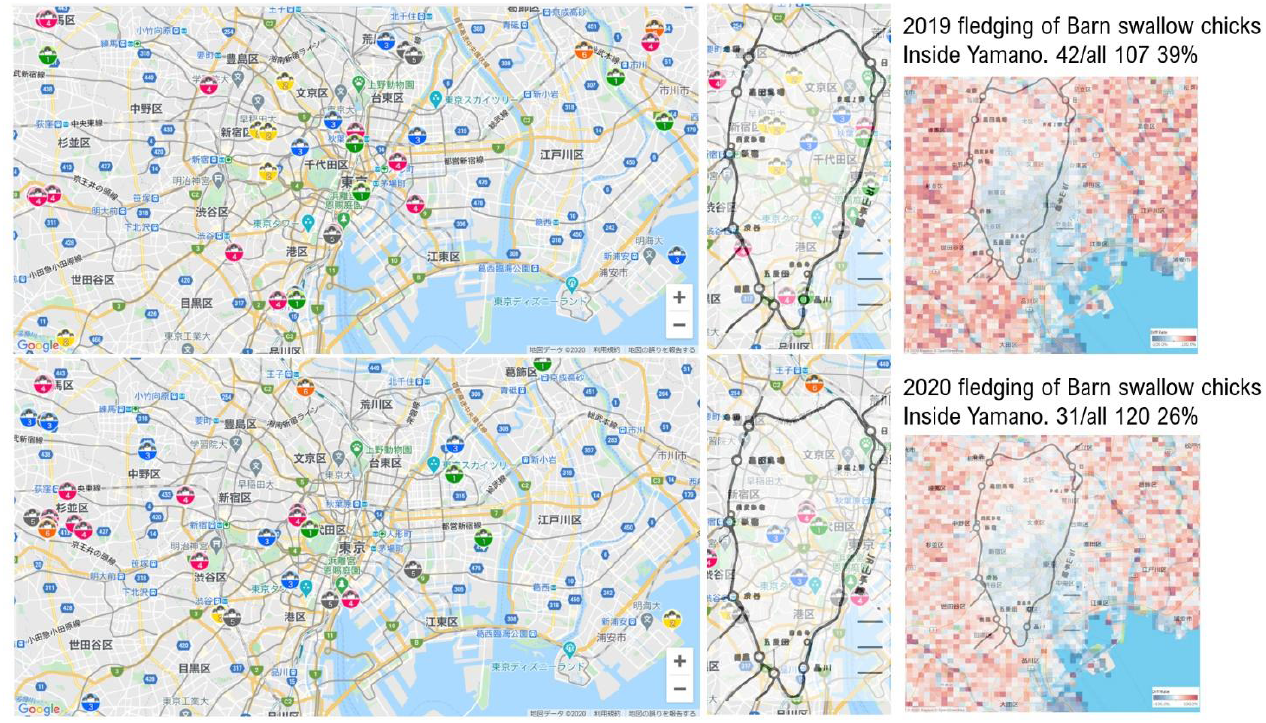
The significant decrease in the number of fledging barn swallow chicks in the area within the Yamanote Line in Tokyo metropolitan in the COVID-19 era. The material^4^ of the National Survey of Barn Swallows of the Wild Bird Society of Japan reveals the number of fledging barn swallow chicks in the entire 23 districts of Tokyo metropolitan and the area within the Yamanote Line in 2019 and 2020. These research results demonstrate that the number of fledging barn swallow chicks in the area within the Yamanote Line in Tokyo metropolitan is extremely low in July 2020, the COVID-19 era, than that in the previous years.

**Figure 3.**
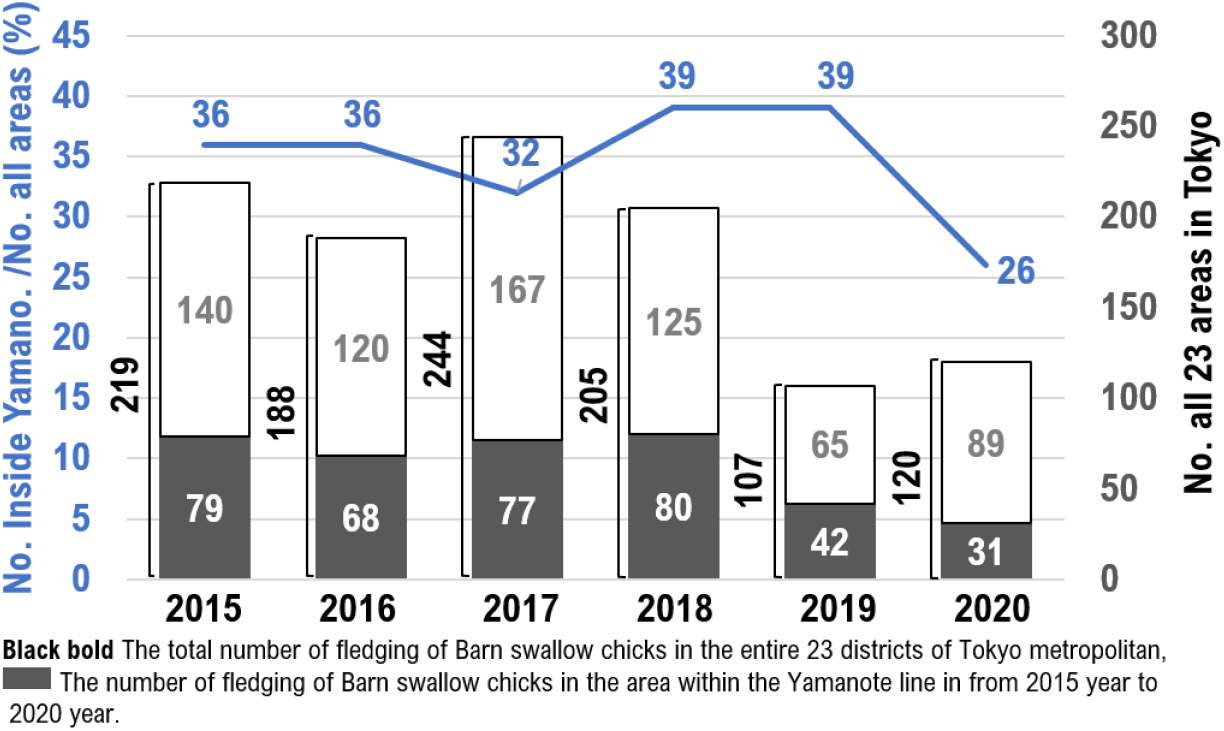
The bar graph shows the number of fledging barn swallow chicks in all 23 districts of Tokyo metropolitan and the area within the Yamanote Line from 2015 to 2020. The line graph shows the ratio of the number of fledging barn swallow chicks in the area within the Yamanote Line to that of fledging swallow chicks in the entire 23 districts of Tokyo metropolitan from 2015 to 2020. These research results demonstrate that the ratio of the number of fledging barn swallow chicks in the area within the Yamanote Line to that of fledging swallow chicks in the entire 23 districts of Tokyo metropolitan is extremely low in July 2020, the COVID-19 era, than that in the previous years.

